# Environmental reservoirs of *Mycobacterium bovis* and *Mycobacterium tuberculosis* in the Ruaha region, Tanzania

**DOI:** 10.1101/790824

**Authors:** Emma R. Travis, Yujiun Hung, David Porter, Goodluck Paul, Robert James, Annette Roug, Midori Kato-Maeda, Rudovick Kazwala, Woutrina A. Smith, Phil Hopewell, Orin Courtenay, Elizabeth M.H. Wellington

## Abstract

This study was designed to investigate the prevalence of members of the Mycobacterium tuberculosis complex (MTBC) in the environment of pastoralists and villagers in the Iringa district, adjacent to the Ruaha National Park in Tanzania. Utilising specific qPCR assays, both *Mycobacterium bovis* and *Mycobacterium tuberculosis* were detected in cattle faeces, boma soil, water and household dust. *M. bovis* was also found in goat faeces and goat boma soil. This is the first report of faecal shedding of *M. bovis* in goats and the first molecular survey of faecal shedding in cattle. The prevalence of both bacterial species varied by village, area, season and sample type. Geographical and temporal correlations across sample types were suggestive of cross species transmission. This non-invasive test has previously been rigorously validated for screening other mammals; in this study it has successfully been applied to detect *M. bovis* and *M. tuberculosis* in livestock faeces and the environment.

## INTRODUCTION

Members of the *Mycobacterium tuberculosis* complex (MTBC) are responsible for transmissible disease in humans, livestock and wildlife. The bacteria within the MTBC have a broad host range; however there are differences in host susceptibility and the specific pathophysiology of the presenting disease, reviewed by (1). *Mycobacterium tuberculosis* (Mtb) is the main cause of human tuberculosis and *Mycobacterium bovis* (Mb) the primary cause of bovine tuberculosis in cattle, other livestock and wildlife. Mb is a re-emerging zoonotic bacterial agent that is of particular concern in sub-Saharan Africa. With an increased recognition of the public health significance of zoonotic TB, the World Health Organization (WHO) and The International Union Against Tuberculosis and Lung Disease convened a meeting in April 2016 and developed a key Zoonotic TB strategy to combat the disease. They published a multisectoral roadmap in 2017 detailing priorities for addressing disease caused by Mb: zoonotic tuberculosis in people and bovine tuberculosis in animals (2). They prioritised the need to improve the scientific evidence base, reduce transmission at the animal-human interface and strengthen intersectoral and collaborative approaches. For zoonotic TB to be successfully addressed it is necessary to consider the burden of the disease in the animal reservoirs and also to consider the risk pathways enabling transmission. Therefore a One Health approach is required which embraces the animal, human and environmental health sectors collectively (3).

Tuberculosis leads to a significant health burden worldwide, with an estimated 10.0 million people suffering from the disease in 2017 and 1.6 million attributed deaths (4). The extent of zoonotic TB is considered to be under-represented due to the similarities in the presentation of the disease. Mb is the main causative agent of zoonotic TB in humans worldwide (reviewed by Muller (5)) with *M. caprae* also responsible for an unknown proportion of incidences. In regions outside Africa, the percentage of TB that is zoonotic is <1.4%, rising to a median of 2.8% in Africa (2); however, there is a disproportional abundance of zoonotic Tb in particular groups and settings. For example, high prevalence of TB attributed to Mb have been reported in Tanzania (16%)(6), Ethiopia (16.3% from human specimen culture)(7), and Zambia (17.9%)(8). HIV is considered an important risk factor for contracting Mb with co-infection associated with increased morbidity and mortality (9). Whilst it is the transmission of Mb from livestock to humans that is the primary concern, there is also increasing evidence of transmission of Mtb from humans to cattle (10-13) in Western Europe as well as across Africa. There are also reports of Mtb infection in wildlife. Whilst most reports are on animals held in captivity (14), Mtb has also been detected in free-ranging wildlife, for example, in banded mongooses *(Mungos mungo)* in Botswana and suricates *(Suricata suricatta)* in South Africa (15). Therefore, it is possible that Mtb infections from humans could negatively impact the health of livestock and also wildlife populations. Furthermore, there is a possibility that livestock and wildlife could represent a Mtb reservoir for reinfection of humans. Whether environmental reservoirs act as a route of disease transmission is also a key element that needs further research. Previous studies revealed that environmental contamination, formed by faecal shedding associated with infection provided a potential and indirect route for transmission of Mb infection (16, 17). The importance of a One Health approach is key; encompassing humans, livestock, wildlife and environmental understanding (Figure 1).

**Figure 1.**
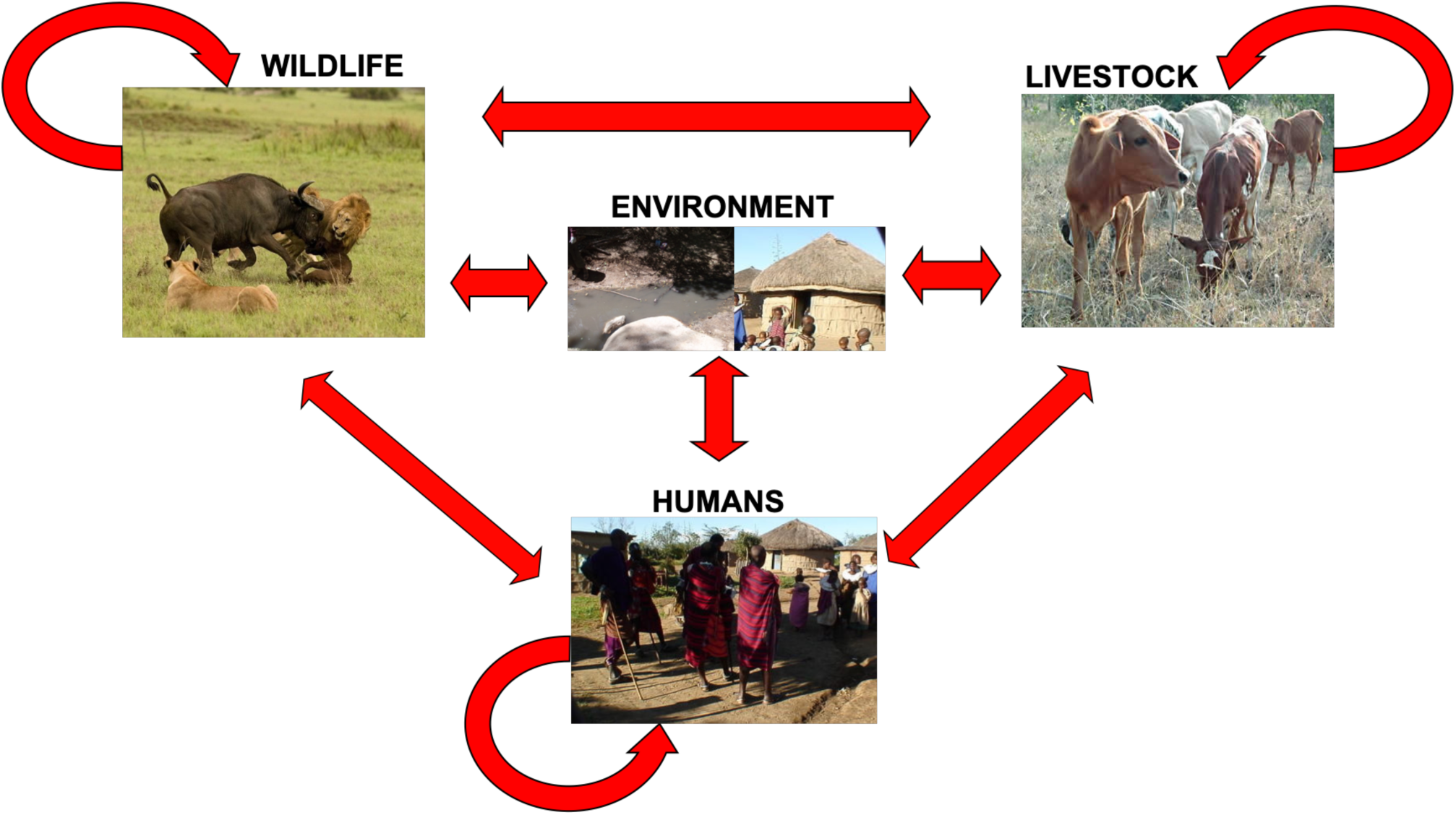
Potential transmission routes for Mb and Mtb. Direct transmission and indirect transmission through the environment are possible.

Developing countries have a multitude of factors that exacerbate the emergence of zoonoses. It is therefore in these locations where the re-emergence of zoonotic disease is having the biggest impact on human health. High rates of human population growth, and resulting land-use change, have culminated in the increased co-habitation of livestock and wildlife. With limited sources of water, livestock and wildlife often converge at these spatially heterogeneous resources which, in turn, increases the risk of zoonotic transmission events(18). These factors have changed the ecological balance between pathogens, animals and humans which has resulted in the emergence and re-emergence of zoonoses. Availability of natural resources in developing countries is key to people’s way of life, with water being the most important of these resources. The density of both humans and animals increase around limited water sources as populations grow. As these water sources are being utilised by both humans and animals for drinking and bathing, they present as key potential reservoirs of infection. Poor animal husbandry practises and sanitation can also increase chances of transmission through faecal contamination of food (19).

The Ruaha ecosystem in Tanzania is one such area witnessing the pressures of an increasingly water-scarce rural environment with humans, livestock and wildlife living in close proximity to each other. The Ruaha ecosystem is one of Tanzania’s largest wild areas which encompasses over 45,000km^2^ and is traversed by the Great Ruaha River and Ruaha National Park. With a high diversity of wildlife and 80% of the human population categorised as rural, both humans and animals rely on natural resources such as water to survive. In the dry season water scarcity increases, with even the Great Ruaha River having no surface water in parts due to recent activities of humans extracting water upstream(20). Previous studies in the Ruaha region have explored the prevalence of MTBC in human, livestock and wildlife hosts(21-25). *Mb* was detected in eight wildlife species inhabiting protected regions on the border of livestock enclosed areas (22). The current study focused on pastoralist communities in the Iringa administrative region, in South-Central Tanzania, near the Ruaha National Park. This area was chosen due to increasing water scarcity, heightened risk of zoonotic transmission via shared resources, and the resultant escalation of human-wildlife conflict in settlements adjacent to the Ruaha park.

This study aimed to establish the prevalence of tuberculous relevant pathogens, namely Mb and Mtb, the environment of pastoralists in the Ruaha area of Tanzania. Furthermore, we aimed to elucidate the presence of Mb and Mtb in the environment within the villages using a non-invasive, sensitive and accurate assay demonstrating shedding of live cells (26-28). In addition, we aimed to explore the influence of season (dry or wet) on prevalence of Mb and Mtb. To achieve these aims, we examined environmental samples for the presence of both Mtb and Mb focusing on cattle and goat faeces, material from the cattle and goat bomas, household dust as well as water from the main water sources in the area. The directionality of transmission is difficult to establish by these methods, instead it is the monitoring of pathogen shedding in the environment which will be studied.

## Materials and Methods

### Study area

The Iringa region was selected as a study site, with pastoralist villages, rivers and water sources. The villages and water sources fell into three regions, that were geographically separated (Figures 1, 2). The regions were populated with Maasai who had similar styles of livestock-rearing typical of pastoralist communities. The regions were regularly monitored by members of the local sampling team who were adept at establishing good community relations. Permissions from all villagers involved were obtained for all sampling. Villagers raised short-horn Zebu (*Bos indicus*) cattle and the majority also kept goat (*Capra hircus*) herds of 20-50 animals with their kids. Cattle and goats were kept in a livestock enclosure (boma) overnight, surrounded by makeshift fences, sticks and vegetation to deter predation. During the day they grazed in a free-range manner with communal grazing grounds and watering points.

**Figure 2.**
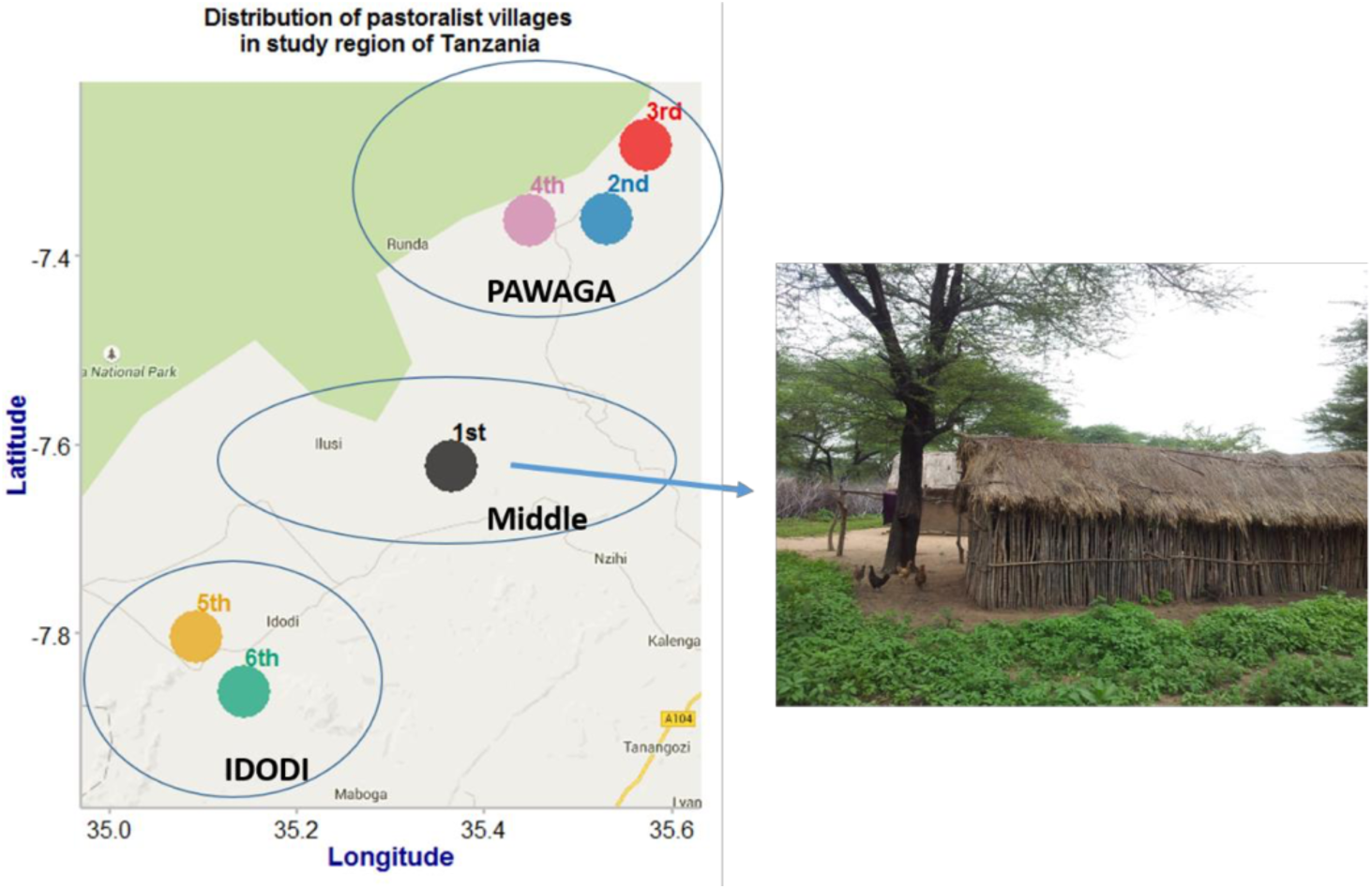
Distribution of six pastoralist villages in the study region of Tanzania. The green area indicates the location of Ruaha National Park.

Six pastoralist villages were selected as sample sites, with 1-3 villages in each region, each village had five or more households and collectively per village approximately 100 or more cattle. Villages were identified from 1-6 in chronological order of investigation in the wet and dry seasons and coordinates recorded using the Garmin eTrex® 10 handheld GPS device (Garmin (Europe) Ltd, Southampton, UK). Figure 1 shows the location of each village and the spread of sample sites over a region of interface between livestock rearing and wildlife. The distribution of water and sediment sample collection was based on watering holes used by the selected villages and chosen from the main or branch river Ruaha (Figure 2). One of the eight places sampled in the wet season was unavailable in the dry season due to water scarcity.

### Sample selection and collection

Samples were taken in September (2012) during the dry and during February (2014) in the wet season. Household samples were taken from the living area of the huts of those who chose to participate in the study, which typically had mud floors. Dust was collected with sterilised trowels into plastic bags from each household and pooled immediately to make a mixed sample to represent individual household. The bag was shaken and mixed thoroughly, prior to three biological replicates of 5 g being collected in Eppendorf tubes.

The top 1 cm^3^ of dried faeces exposed to the sun was removed prior to sample collection to reduce the effect of UV irradiation. At each site, depending on number of livestock and size of livestock enclosure, ten faecal samples were collected from cattle faeces from within the cattle boma and four from goat faeces from within the goat boma. Animals were segregated and faecal samples visually inspected to ensure correct samples were taken. Approximately 2 g faeces were collected using sterilised spatula and placed into a 1.5ml Eppendorf then stored in cooled boxes maintained at 4 °C with a frozen ice pack. Boma samples were taken from the area within the livestock enclosure. The top 3 cm^2^ of dust and vegetation were removed prior to sample collection to reduce the effect of UV radiation. An average of nine samples per boma were collected using the sterilised spatula into the plastic sample bags and pooled immediately to make a mixed sample to represent individual homestead. The bag was shaken prior to subsampling then three biological replicates of 2g each were taken and stored in Eppendorf tubes following the procedure described above.

Water samples were collected from the surface of running river water facing upstream into the current. A total of > 500 ml was collected from each sampling location then each 250 ml water was filtered using 100 ml sterilised plastic syringe and MicrofilV filtration device ([Cat no:] Thermo Fisher Scientific, Leicestershire, UK) and 0.22 μm mixed cellulose esters white gridded filters ([Cat no:] Millipore, MA, USA). Filtration was done *in situ*. After collection the filters were removed from the plastic holder using sterile forceps and air-dried. The filters were then coiled rolled and folded and stored in 2 ml Eppendorf at 4 °C.

Representative mixtures of sediment consisting of sedimentary rock, mud and sand were collected using a sterilised plastic spatula scooped along the bottom of surface river body < 1.5 M deep in the upstream direction. The sediment sample was placed into the 50 ml sterilised plastic universal tube ([Cat no:] Scientific Laboratory Supplies Ltd, Nottingham, UK) and homogenised. Excess water was removed from the container prior to storage.

All samples were stored at 4C prior to being deep frozen using liquid nitrogen then shipped to the UK under Defra licence 51993/194938/3 with appropriate permissions from Tanzanian authorities in a dry shipper under the relevant biohazards labelling UN3373.

The total number of samples analysed in this study was 1513 with sample type and season collected shown in Table 1.

**Table 1.** Samples collected in both wet and dry seasons.

| Sample type | Dry season | Wet season | Total |
| --- | --- | --- | --- |
| Cattle faeces | 394 | 370 | 764 |
| Cattle boma | 90 | 75 | 165 |
| Goat faeces | 116 | 96 | 212 |
| Goat boma | 81 | 64 | 145 |
| Household dust | 90 | 75 | 165 |
| Water from watering hole | 16 | 15 | 31 |
| Sediment from watering hole | 16 | 15 | 31 |
| Total | 803 | 710 | 1513 |

### DNA extraction and qPCR

Mb and Mtb are both Category 3 organisms and as such all testing was conducted at Containment Level 3 at the University of Warwick in line with HSE regulations. Sampling was performed at the University of Warwick to enable it to be performed precisely according to the validated method (27) using an ABI 7500 Fastprep qPCR machine.

Total DNA was extracted from 0.2 g (±0.01 g) of solid samples consisted of faeces, soil, dust and sediment using FastDNA SPIN kit for Soil following manufacture’s instruction. In summary for DNA extraction, 0.2 g sample was added to the lysis matrix tube with 1.4 mm ceramic spheres 0.1 mm silica spheres and 4 mm glass beads which was suitable for soil, sediment and faecal sample as manufacture’s recommendation. Lysis buffer and sodium phosphate buffer were subsequently added into the tube for physical and mechanical disruption using Ribolyser Instrument Precellys 24 (Stretton Scientific Ltd, Stretton, UK). Removal of protein and other materials with protein precipitation buffer was introduced and then performed followed by purification of DNA with ethanol using a centrifugation step was supplied until the end of DNA extraction and collection.

The total number of water samples was 34 filters consisting of 16 from 7 sample sites collected during dry season collection and 15 from 8 sites in the wet season. DNA was extracted from 0.22 μm filter using PowerWater® DNA Isolation Kit following the manufacturer’s instructions. In summary, both chemical and mechanical lysis were employed on cell disruption and removal of protein, cell debris and other organic or any inorganic materials except DNA. The extracts were subjected to high salt solution and then washed with ethanol for purification and consequently DNA was eluted from the membrane using elution buffer prior to storage at −20 °C.

For faecal, boma and household dust samples, total DNA was extracted from samples using 0.2 g (±0.01 g) of solid with FastDNA spin kit for soil (MP Biomedicals) following manufacturer’s instructions. The specific primers and probes for identification of target cells were RD4 deletion region (17) and RD9 (29) with qPCR modified as follows: an initial qPCR of each sample with two technical replicates was conducted using ABI 7500 Fast qPCR machine (ABI) in 96 wells qPCR plate. A first screen determined +/- status of samples by presence of fluorescent signal above threshold using a positive control (10^5^ genome equivalents) and negative control applied in duplicate on each qPCR plate. PCR reactions were set up using 900 nM of each primer (RD4scarF ^5’^TGTGAATTCATACAAGCCGTAGTCG^3’^, RD4scarR ^5’^CCCGTAGCGTTACTGAGAAATTGC^3’^ and RD9F ^5’^TGCGGGCGGACAACTC^3’^, RD9R ^5’^CACTGCGGTCGGCATTG^3’^), 250 nM of Taqman probe (RD4scar probe ^5’^6FAM-AGCGCAACACTCTTGGAGTGGCCTAC—TMR^3’^ and RD9 probe ^5’^6FAM-AGGTTTCA+CCTTCGAC+CC—BBQ^3’^), 1 mg ml^-1^ bovine serum albumen, 12.5 µl of Environmental Mastermix 2.0 (ABI), 10 µl of template DNA and made up to total 25 µl with sterilised molecular grade water (Sigma Aldrich). PCR cycling conditions were 50 °C for 2 min followed by 95 °C for 10 min then 50 cycles of 95 °C for 15 sec and 58 °C for 1 min.

Any positive samples were then analysed by qPCR for enumeration using 3 technical replicates under the same qPCR cycling condition with a full set of reference standards (10^6^ to 1 genome equivalents). If one or two of the technical replicates of the quantification assay exhibited amplification the sample was identified as putative positive for Mb or Mtb depending on primer and probe application and three technical replicates presented amplification the sample was classified as definitely positive for Mb or Mtb. An inhibition control assay previously described was performed to measure the possibility of false negative results due to inhibition (30). Where significant inhibition was detected DNA was re-extracted from frozen aliquots and qPCR assays were repeated.

### Data analysis

Data analysis and spatial studies were performed using R with google map related packages (ggmap, ggplot2, MASS, GLM). Fisher’s exact tests allowed comparisons between prevalences in different seasons or sample types. A generalised linear model was fitted to the count data using a Poisson distribution with the binary data of positivity/negativity of samples modelled with season, sample type and village as categorical factors.

## RESULTS

Sampling took place in the dry season 2012 and the wet season in 2014. Demographic information showed that there were no changes in the population structures of the villages involved between the two dates.

### *M. bovis* detection in cattle faecal and boma samples

Overall prevalence of Mb in the cattle faeces tested was 13.5% (103/764). In the dry season, Mb was detected using the RD4 scar qPCR assay in 17.8% (70/394) of cattle faeces, compared with 8.9% (33/370) Mb positive cattle faeces detected in the wet season (Figure 3). This seasonal variation was significant (Fisher’s exact test two-tailed p=0.0004). In the dry season, Mb was detected in cattle faeces in all 6 of the villages with a prevalence of >= 10%. Village 5 (33.3%) and Village 6 (20.3%) exhibited particularly high levels of prevalence relative to other sample sites. In the wet season, prevalence was lower in all villages apart from Village 6, which showed an increase in prevalence to 30.4%. No cattle faeces from Village 2 tested positive.

**Figure 3.**
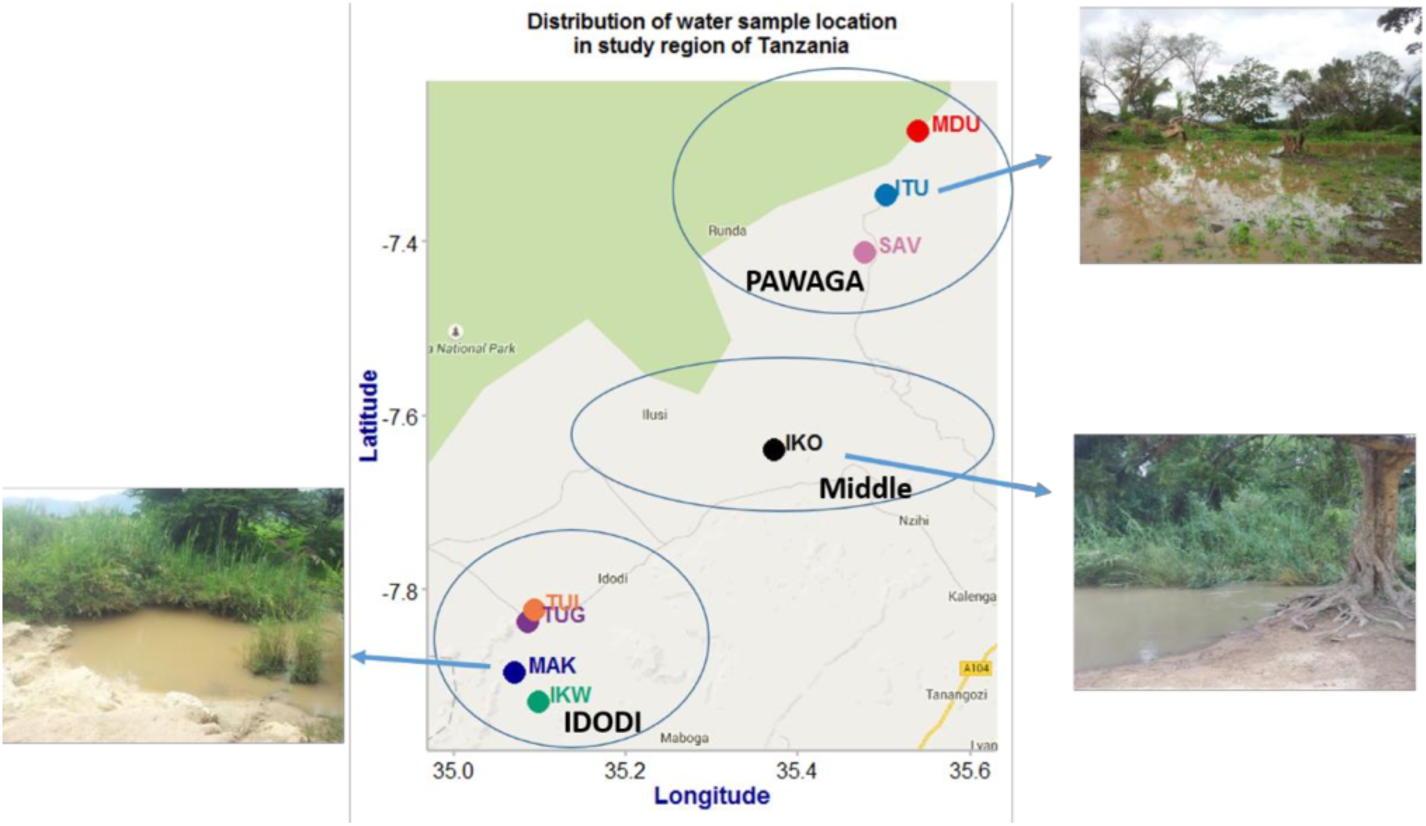
Distribution of water and sediment sample location in the study region of Tanzania. The green area indicates the location of Ruaha National Park

**Figure 3.**
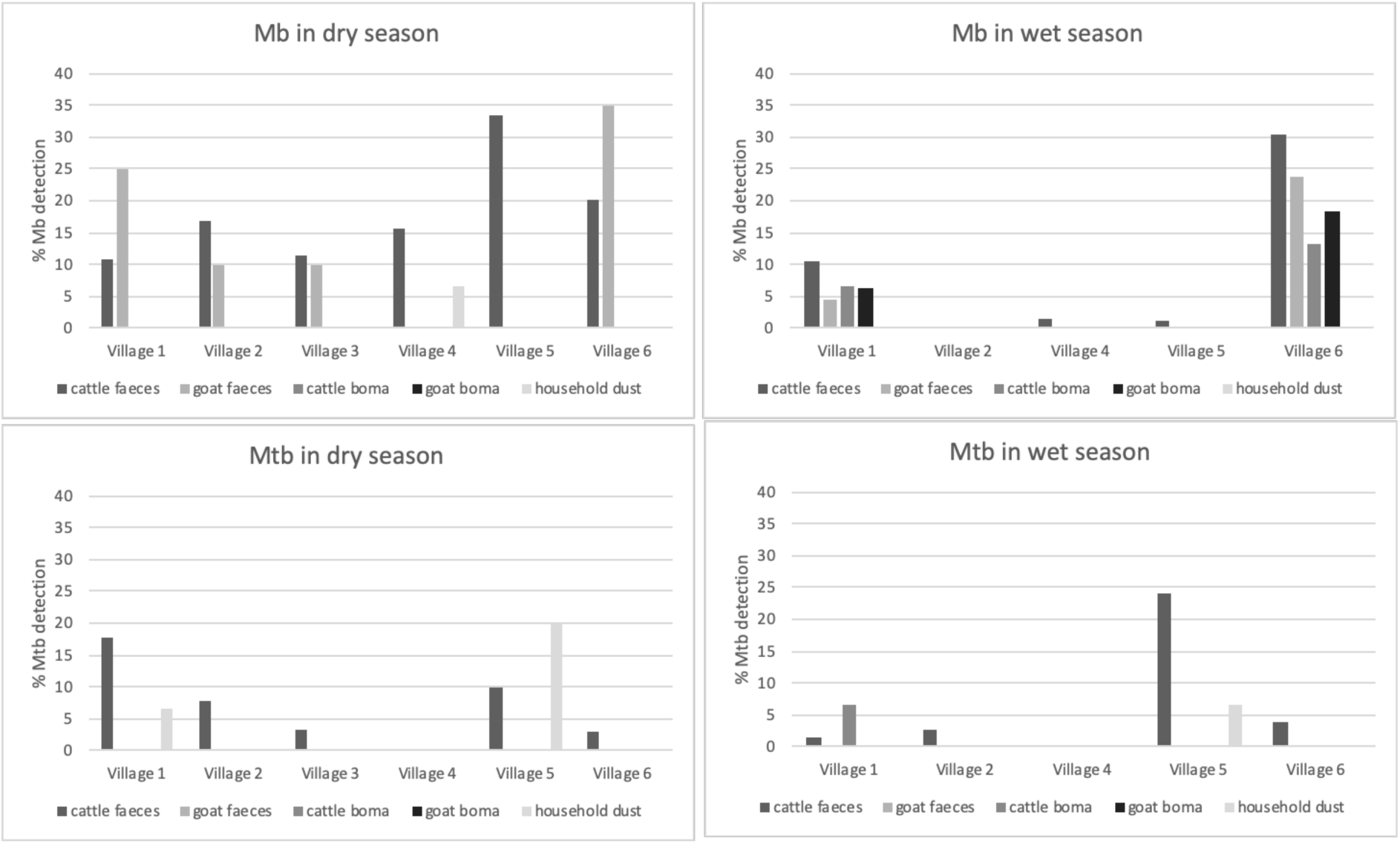
Percentage detection of Mb and Mtb in different sample types across the Villages in dry and wet season. Note it was not possible to sample Village 3 in the wet season due to the inability to access the village.

No cattle boma samples tested positive in the dry season (0/90), whereas in the wet season 4.0% (3/75) samples were positive, although the seasonal difference was not statistically significant (Fisher’s exact test two-tailed p=0.09). In the wet season Village 1 and Village 6 tested positive for Mb in cattle boma samples, these were also the villages that showed the highest prevalence for cattle faeces during this period. All three geographical areas were positive for Mb in the dry season, but area 2 was at significantly lower prevalence for Mb than area 1 and area 3 in the wet season.

### *M. bovis* detection in goat faecal and boma samples

Overall prevalence of Mb in goat faeces was 10.4% (22/212), which, whilst lower than the prevalence in cattle faeces, is not significantly different (Fisher’s exact test two-tailed p>0.05). There was no significant seasonal difference in Mb detection in goat faeces (Fisher’s exact test two-tailed p>0.05) with 13.8% (16/116) samples positive in the dry season compared to 6.3% (6/96) detection in the wet season. No goat boma samples were Mb positive in the dry season (0/81), however, 4.7% (3/64) of goat boma samples were positive in the wet season. This seasonal difference however is not significant (Fisher’s exact test two tailed p=0.08).

In the dry season, Mb was detected in goat faeces from four of the six villages tested, with Villages 1 and 6 with the highest prevalence (25% (5/20) and 35% (7/20) respectively). In the wet season, only Villages 1 and 6 were positive for Mb, with lower prevalences compared with the dry season (4.5% (1/22) and 23.8% (2/21) respectively). Mb was not detected in any goat boma samples collected during the dry season and only Villages 1 and 6 tested positive in the wet season (6.3% (1/16) and 18.2% (2/11) respectively). High prevalences observed in goat samples in Village 1 and 6 correlated with high prevalences in cattle samples.

### *M. tuberculosis* detection in cattle faecal and boma samples

The percentage of cattle faeces that were detected as Mtb positive in the dry season was 7.1% (28/394) which was very similar to that detected in the wet season of 7.0% (26/370) (Fisher’s exact test two tailed p>0.05). One of the cattle was detected as positive with both Mb and Mtb. In cattle boma samples, only one tested positive for Mtb (1.3% (1/75). This sample correlated to a household and village with a Mtb positive cattle faeces. There was a striking difference in Mtb detection between villages across sample types and seasons, with no detection in Village 4, most in Village 5 and high prevalences in Village 1. No Mtb was detected in any of the goat faecal or boma samples in either season.

### Household dust

Across the 30 households tested, prevalence of Mb or Mtb detection in household dust samples was low with 1.1% (1/90) of household dust samples from the dry season testing positive for Mb and 4.4% (4/90) testing positive for Mtb. In the wet season the prevalence was lower, with no Mb detected (0/75) and just 1.3% (1/75) Mtb sample testing positive. Neither the seasonal difference for Mb nor Mtb were significant (Fisher’s exact test two-tailed p>0.05). The sample that tested positive in the wet season was from a household that had also tested positive in the dry season.

Mtb positive household dust samples all came from Village 1 (1/5) and Village 5 (4/5), all samples from different households, with all but one of the Village 5 samples being taken in the dry season. These two villages also have the highest prevalence for Mtb across all other sample types.

The positive Mtb household dust samples were all associated with cattle herds with the highest levels of Mtb. The single Mb positive dust sample originated from a village with Mb positive cattle faeces, indicative of positive herds.

### Water and sediment

Water and sediment samples were taken from watering holes near to the villages. In the dry season, no water or sediment samples were positive for Mb or Mtb (0/16); however, in the wet season, Mb was detected in 26.7% (4/15) water samples but no sediments (0/15). The seasonal difference in water detection was significant (Fisher’s exact test two-tailed p=0.043). This represented samples from all three geographical areas. 6.7% (1/15) of the sediment samples in the wet season tested positive for Mtb, yet no water samples (0/15) tested positive for Mtb. The seasonal variation was not significant for Mtb (Fisher’s exact test two tailed p>0.5). The single Mtb positive water sample was sampled from Area 3, which corresponded with the area containing Villages 5 and 6 which had the highest prevalence of Mtb in both cattle faeces and household dust.

### Statistical correlations

GLM were run with the RD4 dataset to determine the impact of season, village and sampletype on Mb, with all three highly significant factors in Mb prevalence (p<0.001). A similar GLM was run with the RD9 dataset which showed that sample type and village were significant factors in Mtb prevalence in environmental samples (P<0.05), but season was not (p=0.39).

## Discussion

Non-invasive testing of the environment in rural Tanzania has shown that both Mb and Mtb are found across a wide range of environmental samples, raising the possibility of indirect transmission within and between species. Use of qPCR for detecting mycobacterial load in environmental samples has been previously shown to be sensitive and specific when applied to soil, faeces and water (26, 27, 31, 32). There is good evidence that mycobacteria shed into the environment persist and remain viable (33-35). Mb was particularly widespread, with positive samples across all types of environmental samples tested, with varying prevalence according to sample type, season tested and village. Mtb was also widespread, present in all sample types tested, with the exception of goat faeces and goat boma samples. Mtb prevalence also varied according to sample type, season tested and village, with both MTBC members exhibiting spatial and temporal correlations within and across sample types. This study was wide ranging in terms of the environmental samples surveyed, with cattle faeces and cattle boma reflecting infections in cattle, goat faeces and goat boma samples an indication of infection in goats, household dust considered as most likely to derive from human infection and watering hole samples representing a shared facility with potential for cross infection between humans, livestock and wildlife. It is important to consider the implications of this work for each species individually, before considering spatial and temporal correlations and implications for the wider ecosystem.

We detected relatively high prevalence of positive cattle faeces (13.5% overall, although the actual proportion of cattle excreting Mtb or Mb can’t be defined with our approach), with the highest per village prevalence at 33%. There is a considerable body of existing research on Mb in livestock concentrates on cattle herds, the majority of which concentrates on markers of infection, in particular the tuberculin skin test, rather than shedding of the bacterium. In Tanzania, previous studies have shown that Mb in cattle is prevalent, with a study in the Serengeti reporting 14.3% of all cattle tested showing a tuberculin skin test positive result, of which 72.6% of isolates were Mb, with the remainder atypical mycobacterium (23). A study of pastoralist communities in the Ruaha showed 16.5% of herds tested positive to SICCT skin tests (36). In cattle, the presence of Mb in faeces is thought to primarily be derived from swallowing of mucus from the lungs (37) as lesions in the intestine or mesenteric lymph nodes are not common (38). Cassidy (39) observed faecal excretion in cattle with lung lesions but no abdominal lesions. In addition, previous research indicates that within an animal population, some animals are supershedders (40, 41), excreting the pathogen more frequently, at higher concentrations and via more excretion routes. Applying the pareto principle, it can be argued that these few super shedders are responsible for a large proportion of onward transmission. It should also be borne in mind that the majority of studies of cattle excretion have been in developed countries where advanced cases of disseminated tuberculosis in cattle are rare (42) due to regular testing and removal of infected animals. In developing countries with limited or no Mb control measures, the situation may be different with disease being more progressive and outbreaks less managed. In addition, it should be borne in mind that previous studies investigating excretion of Mb in cattle faeces are limited and rely on low sensitivity culture-based techniques. In the current study, it is the excretion of the bacteria via faeces into the environment that is being monitored, using a highly specific molecular approach - qPCR.

Mtb was detected in cattle faeces, with 7.1% testing positive, an unexpected result but Mtb has also been reported in cattle from across the globe (43), with direct transmission from human to cattle linked by strain typing (12). Mtb in cattle have been studied in areas with high incidences of human TB, in particular in Asia and Africa, and reported varying prevalence of Mtb in cattle with 7.4% in Sudan (44), 6.2% in Algeria (45). Reports of infection in Ethiopia indicated the proportion of mycobacterial isolates from infected cattle that were Mtb positive, with findings of 5% (46) and 27% (10). It was noted that prevalence of Mtb was much higher in cattle from herds raised in pastoralist communities, such as those in this study, rather than intensive rearing (10). In Nigeria typing has allowed investigation into origins of the disease, with the cattle strain type being more closely related to those observed in neighbouring Cameroon than circulating in the local human population (47). Mb in wild-ranging animals has also been tested (41, 48).

Mb was detected in 10.4% of goat faeces tested, whereas Mtb was not determined in any samples. Whilst cattle are known to be a maintenance host for Mb, the situation for goats is less clear, with this species only considered a maintenance host in some circumstances. In most cases they are considered as a spill over or dead end host, with infection primarily coming from association with infected maintenance hosts (49). Common grazing of ruminants and infected herds of cattle have been associated with higher tuberculosis infection rates (50) with molecular typing suggesting within farm transmission between cattle and goats (51). We observed that the goats and cattle shared common grazing ground and, whilst a clear correlation exists between cattle and goat sample Mb prevalences within villages, further molecular typing would be required to begin to infer transmission dynamics. The prevalence of Mb infection in goats in rural Africa has been reported to be 4.5% in a study in Nigeria, which quantified detection through post slaughter cattle necropsies and the identification of lesions associated with tuberculosis (52). No previous publications are known to the authors in which shedding of *M. bovis* in goat faeces is described. A molecular study showed spoligotypes of Mb found in goats were also found in humans, suggesting goat to human zoonotic transmission as a possibility (53).

Both Mb and Mtb were detected in household dust samples. These were considered to act as a proxy for human infection, although given the close relationship of the pastoralists and their animals, with goats observed running in and out of households and playing with the children, it cannot be conclusively separated into environmental contamination from humans rather than from the animals they associate with. Houses were observed to be clean with swept mud floors. Families did spit, though this tended to be outside. In part due to distance from the clinics, the pastoralists were unlikely to present at clinics for TB screening so the TB status of the members of the households was unknown. It was particularly notable that the villages in which household dusts were positive for Mtb were also those with the highest levels of Mtb detection in cattle faeces, and where a cattle boma sample was also positive. Whilst this study shows correlation, future studies using molecular typing and whole genome analysis could help reveal disease transmission routes, as has been attempted in Nigeria (47). It would also be useful to investigate in conjunction with strain typing from mycobacteria of human origin.

We explored both temporal and spatial variations in MTBC detection. Firstly, exploring the temporal variations observed; samples were taken during the dry and the wet season. There were some seasonal variations in Mb prevalence within sample types, some of which were significant, whereas there were no significant seasonal differences observed in Mtb prevalence for any sample type. Whilst the highest prevalence of Mb were observed in cattle and goat faeces during the dry season, in contrast, boma samples tested positive during the wet season, but not from samples taken during the dry period. Faecal samples were taken from fresh samples and boma samples were from just below the surface of the livestock enclosure, this discrepancy is likely due to the degradation of mycobacteria via desiccation and UV / white light exposure, despite sampling precautions (sampling from below the surface) to limit this effect (54). In the dry season all animals, wild and livestock use the same watering holes, increasing likelihood of disease transmission and shedding, therefore the watering holes were considered an important possible route of transmission particularly at this time. This, however, was the opposite of what was determined, with Mb and Mtb only being detected in the wet season, consistent with previous research that showed the presence of *M. bovis* present in water samples in Ethiopia (18). Many other factors can also come into play. It was observed that levels in the watering holes were lower than had been expected, perhaps due to decreased water availability from the ground water as a result of removal of water from the upriver areas. In the dry season the impact of UV and white light on mycobacterial viability would be higher. A more intense sampling effort, or larger samples of water would be of interest to determine the extent of shedding across the seasons into this important environmental reservoir with a high potential for disease transmission. The seasonal effect in this study was limited to two sampling occasions with a two year sampling gap between sampling periods. Further sampling occasions would be needed to more clearly differentiate between seasonal differences and changes due to herd movements or natural variation over time.

Secondly, considering the spatial variations in Mb and Mtb prevalence; high prevalences of both MTBC species tested revealed hotspots observed across multiple sample types in the same sampling period, although strain typing would be needed to confirm this. In the current study, there were strong correlations of environmental contaminants from multiple species, with correlations between the prevalence in cattle and goats across the villages, raising the possibility of interspecies transmission. In addition, the positive household dust samples correlated with Mb and Mtb detection respectively in the herds. Presuming that the faecal positive samples indicate infection in the species tested and that the household dust reflects primarily contamination from human sources, the high prevalence across sample types in the same location raises the possibility of transmission between species. The only water sediment sample testing positive for Mtb was from the same area as the villages with the highest prevalence in cattle faeces and household dust, raising the possibility of an environmental transmission route, with intake of food and water considered high risk for indirect transmission (55, 56). Strain typing would allow clearer insights into transmission routes; however, a greater environmental prevalence could be responsible for transmission both between and within species, representing potential transmission between livestock, humans and wildlife. Animals that are super shedders, shedding higher concentrations via more excretion routes, more frequently may be responsible for high levels of transmission (40, 41).

To conclude, non-invasive environmental screening for MTBC has proven a useful tool to enable a targeted investigation of the environment, focussing on faecal samples, livestock enclosure samples, household dust, and water. Spatial and temporal variations were observed in MTBC prevalence, as well as differences between sample type. In Tanzania, there is currently no screening programme for infected animals or culling thereof, allowing disease progression within the animals, increasing the likelihood of shedding, and therefore a higher risk of disease transmission. Whilst detecting Mb in faeces will not detect all infected animals, only those that are shedding, it could be argued that the animals most responsible for the majority of transmission are those which are shedding into the environment. Whilst this study primarily focussed on detection in rural villages, it could be expanded to explore wildlife and utilised further as a tool in trying to understand zoonotic transmission between livestock species, to and from humans as well as into wildlife. In addition to detection of bacterial presence, it could also be expanded to strain typing which would also allow insights into transmission. A recent development in the Wellington laboratory is the use of long read sequencing (nanopore) for MTBC strain typing which could prove a powerful technique to expand this non-invasive environmental screening to allow further insights into within and between species transmission routes (57).

## Acknowledgements

This project was part of the NIH International Collaborations in Infectious Disease Research (ICIDR) U01. RFA-AI-09-010.

